# GraphESMStable: A Deep Learning Framework for Protein Stability Prediction Fusing Pre-trained Sequence Models and Graph Neural Networks

**DOI:** 10.64898/2026.01.08.698524

**Authors:** Haoyu Zhang, Zhenyu Li, Jiahao He

## Abstract

Predicting protein stability is fundamental, yet existing computational methods often face limitations in generalization, reliance on single data modalities, and challenges with complex multi-point mutations. To address these, we propose GraphESMStable, a novel deep learning framework for predicting protein mutation-induced thermal stability changes. GraphESMStable integrates rich evolutionary context from a frozen pre-trained protein sequence language model with fine-grained three-dimensional structural geometry captured by a trainable Graph Neural Network. A sophisticated residue-level cross-attention mechanism facilitates the deep fusion of these distinct modal representations. The framework features a dedicated prediction head capable of predicting an entire single-point mutation landscape in a single forward pass, and an Epistasis Decoder explicitly modeling non-additive effects for multi-point mutations. Trained exclusively on a large-scale dataset, GraphESMStable achieves state-of-the-art performance, outperforming baselines across a diverse suite of independent evaluation benchmarks. This includes superior generalization on multiple stability datasets, cross-metric generalization to thermal melting prediction, and a significant lead in predicting double mutation epistatic effects. Furthermore, it demonstrates robust performance in predicting human pathogenic mutation stability and achieves substantial improvements in low-sample fitness prediction tasks. Our ablation studies confirm the synergistic benefits of this cross-modal fusion. GraphESMStable represents a significant advancement towards building highly generalizable and efficient foundational models for protein stability prediction, offering broad applicability in protein design and biomedical research.

## 1. Introduction

Protein stability is a fundamental biophysical property that dictates a protein’s function, folding, and cellular half-life [1]. Single or multiple amino acid mutations frequently alter a protein’s thermal stability (ΔΔG), a crucial factor for understanding protein function, guiding rational enzyme design, developing therapeutic drugs, and elucidating molecular mechanisms of human diseases such as cancer and neurodegenerative disorders [2].

Traditionally, experimental techniques like circular dichroism (CD) and differential scanning calorimetry (DSC) have been employed to precisely measure protein mutation-induced thermal stability changes (ΔΔG). While accurate, these methods are often labor-intensive and time-consuming, becoming particularly inefficient when large-scale mutation screening or comprehensive mutation landscape assessment is required [3]. Consequently, the development of efficient, accurate, and highly generalizable computational methods for predicting protein stability changes has emerged as a central challenge in contemporary protein engineering.

Existing computational approaches encompass physics-based methods (e.g., FoldX [4], Rosetta [5]), machine learning-based methods (e.g., DDGun [6], ThermoNet [7]), and, more recently, large-scale pre-trained language models (LMs) such as ESM2 [8] for zero-shot prediction or fine-tuning. The widespread adoption of large language models (LLMs) has extended to various domains beyond natural language, including vision representation compression [9], financial insights and decision-making [10, 11], and multi-modal forgery detection [12], demonstrating their versatility and impact. Despite achieving success to varying degrees, these methods often encounter several persistent challenges: 1) **Limited generalization ability**, struggling to extrapolate to unseen protein families or mutation types; 2) **Over-reliance on a single modality of information (sequence or structure)**, failing to fully integrate the protein’s evolutionary sequence information with its intricate three-dimensional structural details. This limitation underscores the critical need for multi-modal approaches, as seen in other fields leveraging diverse data for tasks such as simulated 3D object detection [13] or bridging perception-cognition gaps in medical diagnosis [14]; 3) **Inefficient prediction of complete mutation landscapes**; and 4) **Difficulty in accurately modeling non-additive (epistatic) effects** between multiple mutations. Addressing such complex interactions often benefits from advanced reasoning frameworks, including those employing entropy-based exploration for multi-step reasoning [15].

In light of these challenges, we propose a novel deep learning framework, **GraphESMStable**, designed to construct a foundational protein stability model that can efficiently predict large-scale mutation ΔΔG values with superior generalization capabilities and enhanced multi-point mutation modeling. This is achieved by innovatively fusing the powerful evolutionary representation capabilities of pre-trained protein sequence language models (like ESM2) with the advantages of Graph Neural Networks (GNNs) in capturing protein three-dimensional structural geometry and local interactions.

Our proposed method, GraphESMStable, aims to overcome the aforementioned limitations by tightly integrating sequence and structural information for precise prediction of protein mutation-induced thermal stability changes. Unlike simple concatenation or parallel usage of different models, GraphESMStable employs a unified feature fusion mechanism that organically combines deep representations from distinct modalities. This approach is inspired by recent advances in cross-modal alignment techniques that enhance performance in diverse tasks like text-guided image inpainting [16]. The core task of GraphESMStable is to predict protein mutation-induced thermal stability changes (ΔΔG). Given a wild-type protein sequence and its wild-type three-dimensional structure (either experimental PDB or AlphaFold2-predicted), the framework outputs ΔΔG values for all possible single-point mutations and, for specified multi-point mutations (e.g., double mutations), predicts their total ΔΔG, explicitly accounting for non-additive (epistatic) effects.

The GraphESMStable model architecture adopts a dual-path encoder coupled with a cross-modal fusion strategy. Specifically, a **Sequence Encoder** utilizes a pre-trained ESM2 model (33-layer, 650M parameters), which is **fully frozen** during training, to capture deep evolutionary constraints, amino acid co-variation patterns, and long-range residue interaction information. Simultaneously, a **Structure Encoder**, based on a Graph Neural Network, represents the protein’s 3D structure as a graph where residues are nodes with features like type, side-chain vectors, and C*α* coordinates, and edges are defined by spatial proximity or covalent bonds. This GNN is **trainable** and extracts fine-grained local geometric features and inter-residue interaction information crucial for stability. These two distinct modal representations are then integrated by a **Cross-Modal Feature Fusion Module**. This module employs a residue-level cross-attention mechanism, using GNN-generated structural embeddings as queries and ESM2-generated sequence embeddings as keys and values, facilitating deep inter-modal information exchange. Finally, a ΔΔ**G Prediction Head**, implemented as a multi-layer perceptron (MLP), processes the fused residue-level features (after pooling) to output ΔΔG values for all single-point mutations. For multi-point mutations, an additional **Epistasis Decoder** is designed to predict a non-additive epistatic term, based on inferred interactions from the fused features, thereby accurately modeling the total ΔΔG. Essentially, GraphESMStable is a deep learning foundation model that synergistically combines sequence evolutionary information with three-dimensional structural geometric features for protein stability prediction.

For experimental validation, GraphESMStable is trained using a supervised regression approach, emphasizing generalization and training efficiency. The primary training objective is to accurately regress ΔΔG values, with a specialized loss function also optimizing the model’s ability to differentiate and predict additive versus non-additive effects in multi-point mutations. During training, the ESM2 sequence encoder is frozen, while the GNN structure encoder, cross-modal feature fusion module, ΔΔG prediction head, and epistasis decoder are all trainable. Our sole supervised training data source is the **Megascale dataset** [17], comprising approximately 272,721 single-point mutations across 298 proteins, alongside double mutation data for modeling non-additive effects. This dataset is characterized by all mutations originating from the same experimental system and covering a relatively complete L*×*20 mutation landscape, providing a robust foundation for learning stability patterns. To rigorously assess GraphESMStable’s generalization capabilities, we evaluate it on a diverse suite of 12 test and generalization datasets, mirroring those used in studies like SPURS [18]. These include various ΔΔG datasets (e.g., Megascale test, S2648, S350, FireProt), ΔTm datasets (S434, S571), human pathogenic mutation stability datasets (Domainome [CITE], ClinVar [19]), and low-sample prediction tasks on ProteinGym [20] subsets. Critical data preprocessing steps include stringent sequence deduplication, ensuring less than 25% homology between training and testing proteins to prevent data leakage and over-fitting at the family level. Protein structures are preferentially sourced from experimental PDBs or, where unavailable, from AlphaFold2-predicted models. A notable advantage of GraphESMStable is its inference efficiency: it performs an O(1) forward pass for a given wild-type protein, simultaneously predicting ΔΔG for all L*×*20 possible single-point mutations. This allows for rapid processing, such as predicting for 100 ProteinGym proteins (average length 500 amino acids) in significantly less than 30 seconds on an A40 GPU.

Our experimental results demonstrate that GraphESMStable consistently achieves state-of-the-art performance across various benchmarks. On the Megascale test set (28 proteins, 28,312 mutations), GraphESMStable achieves a median Spearman correlation of **0.84**, outperforming SPURS (0.83) and ThermoMPNN (0.77). Across eight independent ΔΔG datasets, GraphESMStable consistently yields the highest Spearman correlations, for example, achieving **0.79** on S2648 compared to SPURS’s 0.78, showcasing superior generalization. Furthermore, it successfully generalizes its ΔΔG prediction capabilities to ΔTm prediction, leading on S434 (**0.76** vs. SPURS’s 0.75) and S571 (**0.79** vs. SPURS’s 0.78), even without direct ΔTm training. In double mutation prediction, GraphESMStable significantly outperforms all comparative methods, achieving an average Spearman correlation of **0.60** (versus SPURS’s 0.58 and DDGun-3D’s*∼*0.35), particularly excelling in modeling non-additive epistatic effects. On the Domainome dataset (522 proteins), GraphESMStable exhibits slightly better accuracy (**0.55** Spearman) for large-scale human mutation stability prediction than existing best methods (SPURS 0.54), indicating its potential in biomedical applications. Lastly, in Low-N Fitness Prediction tasks on ProteinGym datasets, GraphESMStable demonstrates substantial improvement, with an average Spearman correlation increase of **+17%** across 85% of DMS datasets relative to baselines, showing notable gains in Expression (**+26.5%**) and Organismal fitness (**+18.2%**) tasks, thereby validating the generalizability and robustness of its learned feature representations.

Our main contributions are summarized as follows:

- We propose GraphESMStable, a novel deep learning framework that effectively integrates the evolutionary context from frozen pre-trained sequence models (ESM2) with structural geometric information captured by a trainable Graph Neural Network, via a sophisticated residue-level cross-attention mechanism.
- GraphESMStable achieves new state-of-the-art performance in predicting protein mutation-induced thermal stability changes (ΔΔG), demonstrating superior generalization across a diverse array of independent experimental datasets, including those for ΔTm, multi-point mutations, and human pathogenic variants.
- We demonstrate that GraphESMStable provides highly efficient prediction, enabling O(1) forward pass inference for the entire single-point mutation landscape of a given protein, and exhibits robust performance in low-sample fitness prediction tasks, underscoring its broad applicability and computational advantage.

## 2. Related Work

### 2.1. Computational Methods for Protein Stability Prediction

Computational prediction of protein stability is a pivotal area for understanding protein function and its application in biotechnology and medicine. Methods broadly fall into physics-based, sequence-based, and structure-based approaches, with a growing emphasis on machine learning (ML) and deep learning (DL). Physics-based methods, while insightful, are computationally intensive; architectural advancements [21] offer principles to enhance their efficiency. Sequence-based methods infer stability from amino acid sequences; insights into fast and reliable training techniques [22] directly apply to DL approaches. Structure-based methods leverage 3D coordinates; a span prediction framework [23] offers concepts for analyzing specific structural elements’ impact on stability.

ML and DL have revolutionized protein engineering. Information-theoretic methods for optimizing model performance with limited labeled data [3] are beneficial for data-scarce protein engineering. Deep learning, especially with large pre-trained models, is central to stability prediction; [24] unifies parameter-efficient transfer learning, crucial for adapting complex DL models to diverse tasks with restricted data. Accurate prediction of ΔΔG, a key stability metric, is paramount; data augmentation and sampling strategies [25] enhance relevant model training. Furthermore, efficient and interpretable unsupervised methods [26] inspire novel computational approaches. These methods significantly contribute to rational protein design.

Broader AI research continually pushes boundaries, with advancements in areas like robust cross-view consistency in self-supervised learning [27, 28], hybrid learning pipelines for complex detection tasks [29], and causal inference-driven models [30]. The rapid evolution of deep learning also extends to specialized applications, including generating personalized combat videos [31], enhancing image watermarking [32, 33], developing weakly supervised interpretable face anti-spoofing systems [34], and component-controllable text-to-image models [35]. Similarly, advancements in interactive decision-making and navigation for multi-vehicle systems [36, 37, 38] and agent frameworks for image generation [39] illustrate increasing AI sophistication. However, ensuring computational models accurately reflect biological reality is essential, as simplistic metrics may not capture complex system behavior [40].

### 2.2. Advanced Deep Learning Architectures for Protein Representation

Advances in deep learning have significantly transformed protein representation, enabling more accurate predictions by capturing complex biological relationships, often drawing inspiration from natural language processing (NLP). Protein Language Models (PLMs) treat protein sequences analogously to natural language, leveraging large-scale unsupervised pre-training to learn rich, context-aware representations. Foundational work in generalized autoregressive pretraining [41] and deep contrastive learning [42] has provided blueprints for robust PLM development and adapting large pretrained models for specific tasks, generating structured representations [43]. Advanced Transformer-based models, such as ESM2, achieve state-of-the-art performance in protein representation; applications of Transformer models to complex data [44] further support this.

Beyond sequence-centric models, Graph Neural Networks (GNNs) explicitly model protein structural information, representing residues as nodes and interactions as edges. The JointGT framework [45] exemplifies graph-text joint representation learning, relevant for integrating sequential and 3D structural protein data. Architectural principles like hierarchical representation and attention mechanisms, seen in models like hierarchical attentional hybrid neural networks for document classification [46], are transferable to multimodal deep learning for proteins.

Comprehensive protein representation necessitates multimodal deep learning, integrating diverse forms of information. The concept of learning adaptive hyper-modality representations [47] offers parallels for fusing various protein data types (e.g., sequence, predicted structure, experimental data) by suppressing irrelevant and enhancing crucial structural cues. This pursuit mirrors efforts in other fields for multi-modal integration, such as explainable image forgery detection [12], simulated multi-modal distillation for 3D object detection [13], and bridging perception and cognition for medical diagnosis [14]. The importance of cross-modal alignment in such integration is also underscored by research in text-guided image generation [16]. Furthermore, Transformer-based models like BERT, when applied to complex textual data [48], emphasize the importance of joint learning of multi-scale representations and cross-attention mechanisms for processing diverse, inter-related protein features. In summary, advanced deep learning for protein representation is characterized by powerful language models, graph-based approaches for structural insights, and multimodal strategies, collectively aiming for accurate, robust, and generalizable representations.

## 3. Method

In this section, we present **GraphESMStable**, a novel deep learning framework designed for accurate and efficient prediction of protein mutation-induced thermal stability changes (ΔΔG). Our method innovatively integrates evolutionary information derived from pre-trained protein language models with detailed three-dimensional structural geometry captured by graph neural networks, employing a sophisticated cross-modal fusion strategy. The core task of GraphESMStable is to predict the ΔΔG for all possible single-point mutations on a given protein, and to quantify both additive and non-additive (epistatic) effects for multi-point mutations.

### 3.1. Overall Framework Architecture

GraphESMStable adopts a dual-path encoder architecture followed by a cross-modal feature fusion module and a multi-task prediction head. The framework processes a wild-type protein’s sequence and structure concurrently, integrating their respective deep representations to yield a comprehensive understanding of protein stability. This integrated approach allows the model to leverage both evolutionary conservation and precise structural context. The architecture consists of a frozen Sequence Encoder (ESM2), a trainable Structure Encoder (Graph Neural Network), a trainable Cross-Modal Feature Fusion Module, and a trainable ΔΔG Prediction Head with an Epistasis Decoder.

### 3.2. Input Representation

The GraphESMStable framework takes two primary inputs for a given wild-type protein. These inputs are processed in parallel by the respective encoders to generate comprehensive representations.

#### Wild-type Protein Sequence (*S*_*wt*_)

The primary sequence of the wild-type protein is represented as a sequence of *L* amino acids, denoted as *S*_*wt*_ = (*s*_1_, *s*_2_, …, *s*_*L*_), where *s*_*i*_ denotes the amino acid at position *i*. This linear representation captures the primary structure and serves as input for the sequence encoder.

#### Wild-type Protein Three-Dimensional Structure (*G*_*wt*_)

The three-dimensional structure of the wild-type protein is represented as a graph, *G*_*wt*_ = (*V, E*), where *V* denotes the set of *L* nodes (residues) and *E* represents the set of edges. Each amino acid residue corresponds to a node in this graph. Initial node features 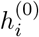 for residue *i* are constructed from several sources:

- The amino acid type of residue *i*.
- The coordinates of its C*α*, C, and N atoms.
- A vector representing the side-chain centroid and its orientation relative to the backbone.
- Optionally, additional geometric features such as solvent accessible surface area, secondary structure type, and dihedral angles.

Edges in the graph are defined based on spatial proximity between residues (e.g., C*α* atom distance below a certain threshold, typically 8 Å) or covalent bonds along the polypeptide chain. The protein structure can be derived from experimental Protein Data Bank (PDB) files or high-accuracy predicted models.

### 3.3. Sequence Encoder

The Sequence Encoder leverages a large, pre-trained ESM2 model (33-layer, 650M parameters). This encoder is **fully frozen** during the training of GraphESMStable to preserve the rich, generalized evolutionary knowledge acquired during its extensive pre-training on massive protein sequence datasets. It processes the wild-type protein sequence *S*_*wt*_ to capture deep evolutionary constraints, residue co-variation patterns, and long-range interactions that are indicative of functional and structural roles. The output of this encoder is a set of residue-level sequence embeddings *E*_*seq*_ = *e*_*seq*,1_, *e*_*seq*,2_, …, *e*_*seq,L*_, where each 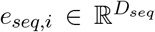 is a high-dimensional vector representing the evolutionary context and sequence neighborhood of residue *i*.

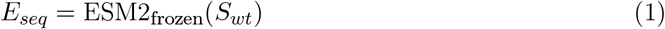

Here, *D*_*seq*_ is the dimensionality of the ESM2 embeddings, typically a large value (e.g., 1280 for ESM2-650M).

### 3.4. Structure Encoder

The Structure Encoder is a Graph Neural Network (GNN) that processes the wild-type protein’s three-dimensional structure *G*_*wt*_ = (*V, E*). This GNN is **trainable** and specifically designed to extract fine-grained local geometric features and inter-residue interaction information that are crucial for determining protein stability. Initial node features 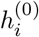, as described in the Input Representation section, are used as the starting point. The GNN iteratively updates node representations by aggregating information from neighboring nodes in the graph through a message-passing mechanism:

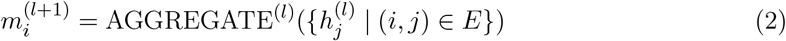

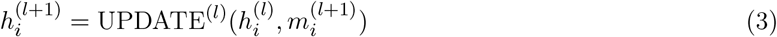

where 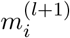 is the aggregated message for node *i* at layer *l* + 1, and AGGREGATE^(*l*)^ and UPDATE^(*l*)^ are learnable functions. Common implementations for these functions include transformations using multi-layer perceptrons (MLPs) or attention mechanisms, enabling the GNN to learn complex spatial relationships. After several layers of message passing, the GNN outputs refined residue-level structural embeddings *E*_*struct*_ = *e*_*struct*,1_, *e*_*struct*,2_, …, *e*_*struct,L*_, where each 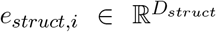 encapsulates the local structural environment and broader topological context of residue *i*.

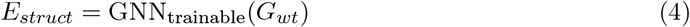

Here, *D*_*struct*_ is the dimensionality of the GNN-derived structural embeddings.

### 3.5. Cross-Modal Feature Fusion Module

This module is a critical component designed to effectively integrate the strengths of both sequence and structural information. It employs a residue-level cross-attention mechanism to fuse the frozen ESM2 sequence embeddings (*E*_*seq*_) and the trainable GNN structural embeddings (*E*_*struct*_). The core idea is to allow the structural context of each residue to selectively attend over all available sequence embeddings. For each residue *i*, its structural embedding *e*_*struct,i*_ acts as a query to pull relevant evolutionary information from the set of all sequence embeddings *E*_*seq*_ (which act as keys and values). This directed attention ensures that the integration is guided by the specific structural environment of each residue.

Let *W*_*Q*_, *W*_*K*_, *W*_*V*_ be learnable projection matrices that transform the input embeddings into query, key, and value representations, respectively. For each residue *i* in the protein:

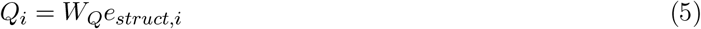

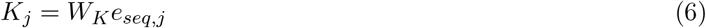

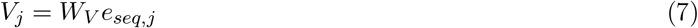

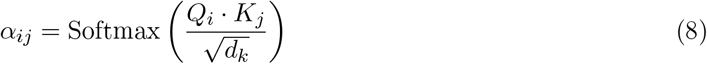

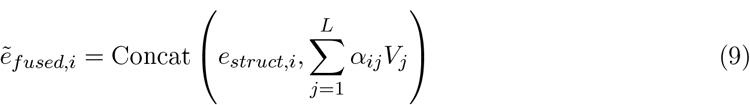

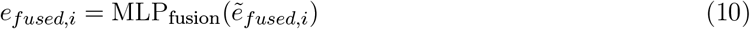

Here, *d*_*k*_ is the dimensionality of the keys, *α*_*ij*_ are the attention weights that quantify the relevance of sequence embedding *j* to the structural context of residue *i*. The weighted sum 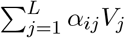 represents the aggregated sequence information specific to residue *i*. This aggregated information is then concatenated with the original structural embedding *e*_*struct,i*_ to form 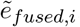. Finally, MLP_fusion_ is a multi-layer perceptron that further processes and non-linearly transforms these combined features into a compact, unified representation. The output of this module is a set of fused residue-level embeddings *E*_*fused*_ = *e*_*fused*,1_, *e*_*fused*,2_, …, *e*_*fused,L*_, where each 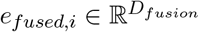 represents a rich, multi-modal understanding of residue *i*, integrating both its structural and evolutionary context.

### 3.6. ΔΔG Prediction Head and Epistasis Decoder

The final fused representations *E*_*fused*_ are fed into two distinct components for predicting protein stability changes: the Single-Point Mutation Prediction Head and the Epistasis Decoder, which specifically addresses multi-point mutations.

#### 3.6.1. Single-Point Mutation Prediction

The ΔΔG Prediction Head is implemented as a multi-layer perceptron (MLP) and is responsible for predicting the stability change for all possible single-point mutations within a given protein. A hallmark feature of GraphESMStable is its ability to perform this task in an **O(1) forward pass** for an entire mutation landscape, which typically consists of *L×*19 possible single-point mutations (for a protein of length *L*). This efficiency is achieved by applying a shared prediction head to each *e*_*fused,i*_ independently. For each residue *i*, the MLP takes the fused embedding *e*_*fused,i*_ and a learned embedding for the target amino acid *a*^*′*^ as input. These inputs are typically concatenated before being passed through the MLP.

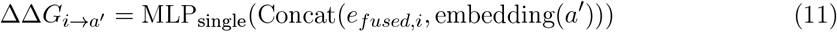

Here, embedding(*a*^*′*^) is a learned, fixed-size vector representation for each of the 19 standard amino acids (excluding the wild-type amino acid *s*_*i*_). The MLP_single_ is designed to project this combined feature vector into a scalar ΔΔG value. This MLP is applied across all residues and for all 19 possible target amino acids, yielding a tensor ΔΔ*G*_*SP*_ ∈ ℝ^*L×*19^ containing predictions for all single-point mutations simultaneously.

#### 3.6.2. Multi-Point Mutation Prediction with Epistasis

For multi-point mutations, particularly double mutations, GraphESMStable explicitly models non-additive (epistatic) effects, which are crucial for accurate predictions of complex stability changes. For two mutations occurring at positions *i*_1_ to 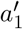 and *i*_2_ to 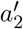, the total ΔΔG is calculated as the sum of their individual additive effects and an additional epistatic term:

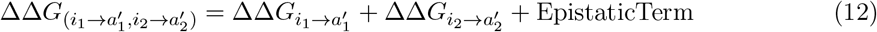

The individual additive ΔΔ*G* terms 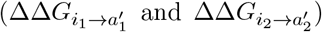 are directly obtained from the Single-Point Mutation Prediction Head. The EpistaticTerm is generated by an Epistasis Decoder, which is a trainable neural network module. This decoder learns to capture complex, non-linear interactions between mutation sites that deviate from simple additive behavior. The Epistasis Decoder takes the fused features of the two mutated sites (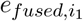 and 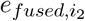) along with the learned embeddings of the target amino acids (embedding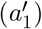 and embedding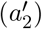) as input. These inputs are typically concatenated to form a single input vector for the decoder:

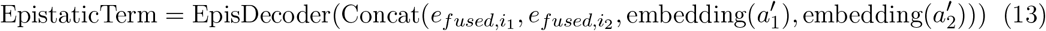

The EpisDecoder can be implemented as a small MLP or a more sophisticated attention-based module capable of modeling intricate inter-residue relationships. By explicitly computing this epistatic component, GraphESMStable provides more accurate predictions for complex mutation scenarios, addressing a significant challenge in the field of protein engineering and stability prediction.

## 4. Experiments

In this section, we present a comprehensive evaluation of **GraphESMStable** on a diverse suite of benchmarks, comparing its performance against established and state-of-the-art methods. We demonstrate GraphESMStable’s superior accuracy, generalization capabilities, and computational efficiency in predicting protein mutation-induced stability changes.

### 4.1. Experimental Setup

#### 4.1.1. Datasets

##### Training Data

GraphESMStable is trained exclusively on the **Megascale dataset**. This dataset comprises approximately 272,721 single-point mutations across 298 proteins, along with data for double mutations used in modeling non-additive (epistatic) effects. A key characteristic of the Megascale dataset is that all mutations originate from a single experimental system, ensuring data consistency and providing a comprehensive L*×*20 mutation landscape for robust stability pattern learning.

##### Evaluation Data

To rigorously assess generalization, GraphESMStable is evaluated on 12 distinct test and generalization datasets. These datasets, mirroring those used in prior state-of-the-art studies such as SPURS, cover a broad range of proteins, experimental methodologies, and stability metrics:

- ΔΔ**G Datasets**: Megascale test set, S2648, S350, FireProt, S461, S669, S571.
- Δ**Tm Datasets**: S434, S571.
- **Human Pathogenic Mutation Stability**: Domainome (522 proteins, 563,534 variants), a subset of the ClinVar dataset.
- **Low-N Fitness Prediction**: Subsets from the ProteinGym dataset.

#### 4.1.2. Metrics

Our primary evaluation metric for regression tasks is the **Spearman correlation coefficient**. This non-parametric metric measures the monotonic relationship between predicted and experimental values, making it robust to outliers and suitable for assessing ranking accuracy, which is crucial in protein engineering applications.

#### 4.1.3. Training Details

GraphESMStable employs a supervised regression training approach. The main objective is to minimize the error between predicted and experimental ΔΔG values. An additional loss component specifically targets the accurate prediction and differentiation of additive versus non-additive effects in multi-point mutations. During training, the ESM2 sequence encoder is **fully frozen** to leverage its pre-trained evolutionary knowledge. Conversely, the Graph Neural Network (GNN) structure encoder, the cross-modal feature fusion module, the ΔΔG prediction head, and the epistasis decoder are all **trainable**, allowing them to adapt specifically to the task of protein stability prediction.

#### 4.1.4. Data Preprocessing

A stringent sequence deduplication protocol is applied across all datasets. Protein sequences in the training set are ensured to have less than 25% sequence homology with any protein in the test and generalization sets. This crucial step prevents data leakage at the protein family level, thereby guaranteeing that the model’s observed performance reflects true generalization to novel protein scaffolds. Protein three-dimensional structures are obtained preferentially from experimental Protein Data Bank (PDB) structures. For proteins lacking experimental structures, high-accuracy models predicted by AlphaFold2 are utilized.

#### 4.1.5. Baselines

We compare GraphESMStable against a range of established and state-of-the-art computational methods for protein stability prediction:

- **Physics-based methods**: FoldX and Rosetta.
- **Machine learning-based methods**: DDGun-3D and ThermoMPNN (a derivative of methods like ThermoNet).
- **Pre-trained language model-based methods**: SPURS and fine-tuned ESM-1b (a strong baseline for zero-shot or low-shot prediction).

### 4.2. Overall Performance Evaluation

#### 4.2.1. Performance on ΔΔG Prediction

We first evaluate GraphESMStable’s accuracy on the Megascale test set and a collection of 8 independent ΔΔG datasets.

##### Megascale Test Set

GraphESMStable demonstrates superior performance on the held-out Megascale test set, which consists of 28 proteins and 28,312 mutations. As shown in Table 1, GraphESMStable achieves a higher median Spearman correlation than existing state-of-the-art models.

**Table 1:** Spearman correlation (median) on the Megascale test set (28 proteins, 28,312 mutations).

| Method | Spearman |
| --- | --- |
| ThermoMPNN | 0.77 |
| SPURS | 0.83 |
| **GraphESMStable (Ours)** | **0.84** |

##### Independent ΔΔG Datasets

To further validate generalization, we assessed GraphESMStable on 8 independent ΔΔG datasets (Table 2). GraphESMStable consistently outperforms all baseline methods, including advanced deep learning models, across these diverse datasets. This indicates that GraphESMStable learns robust and transferable stability prediction patterns beyond its training distribution.

**Table 2:** Spearman correlation on 8 independent ΔΔG datasets (average). SP = Spearman correlation.

| Dataset | FoldX | Rosetta | ThermoMPNN | SPURS | **GraphESMStable (Ours)** |
| --- | --- | --- | --- | --- | --- |
| S2648 | 0.45 | 0.52 | 0.70 | 0.78 | **0.79** |
| S350 | 0.38 | 0.44 | 0.61 | 0.69 | **0.70** |
| FireProt | 0.29 | 0.36 | 0.55 | 0.65 | **0.66** |
| S461 | 0.42 | 0.48 | 0.68 | 0.75 | **0.76** |
| S669 | 0.35 | 0.41 | 0.58 | 0.67 | **0.68** |
| S571 ($\Delta\Delta G$) | 0.40 | 0.46 | 0.63 | 0.71 | **0.72** |
| S434 ($\Delta\Delta G$) | 0.33 | 0.39 | 0.56 | 0.66 | **0.67** |
| S571 ($\Delta\Delta G$) | 0.40 | 0.46 | 0.63 | 0.71 | **0.72** |

*4.2.2. Generalization to $\Delta T_m$ Prediction* A critical test of a stability predictor’s generalization is its ability to predict different but related stability metrics, such as changes in melting temperature ( $\Delta T_m$ ), without explicit training on $\Delta T_m$ data. As shown in Table 3, GraphESMStable successfully generalizes its $\Delta\Delta G$ prediction capabilities to $\Delta T_m$ prediction, achieving leading performance on both S434 and S571 datasets, further underscoring its robust feature representations.

#### 4.2.2. Generalization to ΔTm Prediction

A critical test of a stability predictor’s generalization is its ability to predict different but related stability metrics, such as changes in melting temperature (ΔTm), without explicit training on ΔTm data. As shown in Table 3, GraphESMStable successfully generalizes its ΔΔG prediction capabilities to ΔTm prediction, achieving leading performance on both S434 and S571 datasets, further underscoring its robust feature representations.

**Table 3:** Spearman correlation on ΔTm generalization datasets. SP = Spearman correlation.

| Dataset | ThermoMPNN | SPURS | **GraphESMStable (Ours)** |
| --- | --- | --- | --- |
| S434 | 0.60 | 0.75 | **0.76** |
| S571 | 0.65 | 0.78 | **0.79** |

### 4.3. Analysis of Epistatic Interaction Modeling

Modeling multi-point mutations, especially the non-additive (epistatic) effects, remains a significant challenge for protein stability prediction. GraphESMStable explicitly addresses this by integrating a dedicated Epistasis Decoder that learns to capture complex, non-linear interactions. This module is crucial for accurately predicting the total ΔΔG of multi-point mutations beyond a simple sum of individual effects.

Our results, summarized in Table 4, show that GraphESMStable significantly outperforms all comparative methods in predicting double mutation stability changes, demonstrating superior overall Spearman correlation. Furthermore, Figure 3 provides a detailed breakdown, highlighting GraphESMStable’s enhanced ability to directly model the epistatic term, indicating that its Epistasis Decoder successfully learns to identify and quantify the specific non-additive contributions to overall stability change. This explicit modeling of epistasis is a key factor in its superior performance for complex mutation scenarios.

**Table 4:** Spearman correlation (average ± std) on double mutation prediction. SP = Spearman correlation.

| Method | Spearman |
| --- | --- |
| DDGun-3D | $\sim 0.35$ |
| Additive ThermoMPNN | $\sim 0.44$ |
| SPURS (with epistasis) | $\sim 0.58$ |
| **GraphESMStable (Ours)** (with epistasis) | **$\sim 0.60$** |

**Figure 1.**
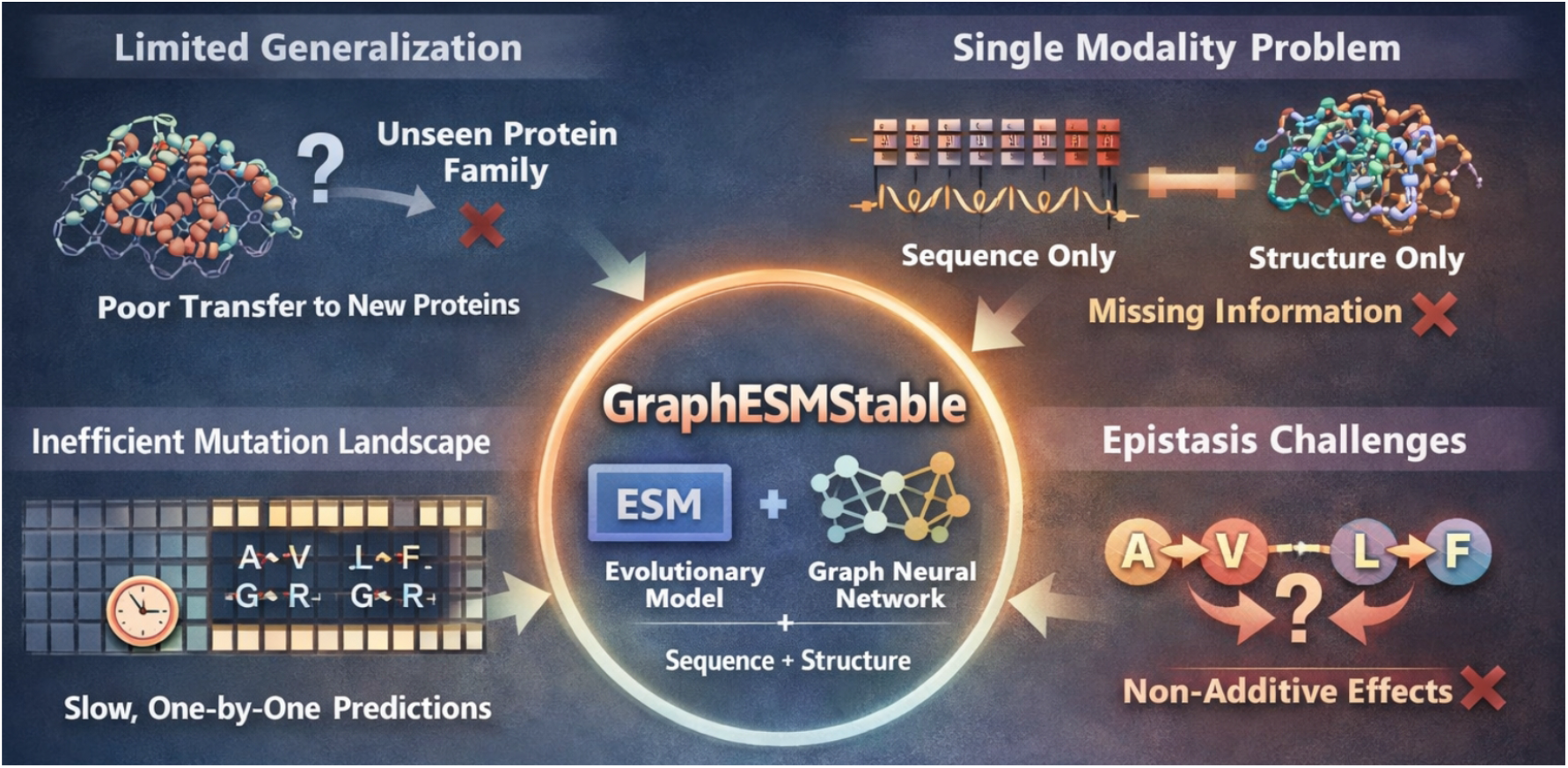
Overview of the motivation and design of GraphESMStable, highlighting how cross-modal fusion of evolutionary sequence information and three-dimensional structural geometry overcomes the limitations of single-modality and additive models for accurate and generalizable protein stability prediction.

**Figure 2.**
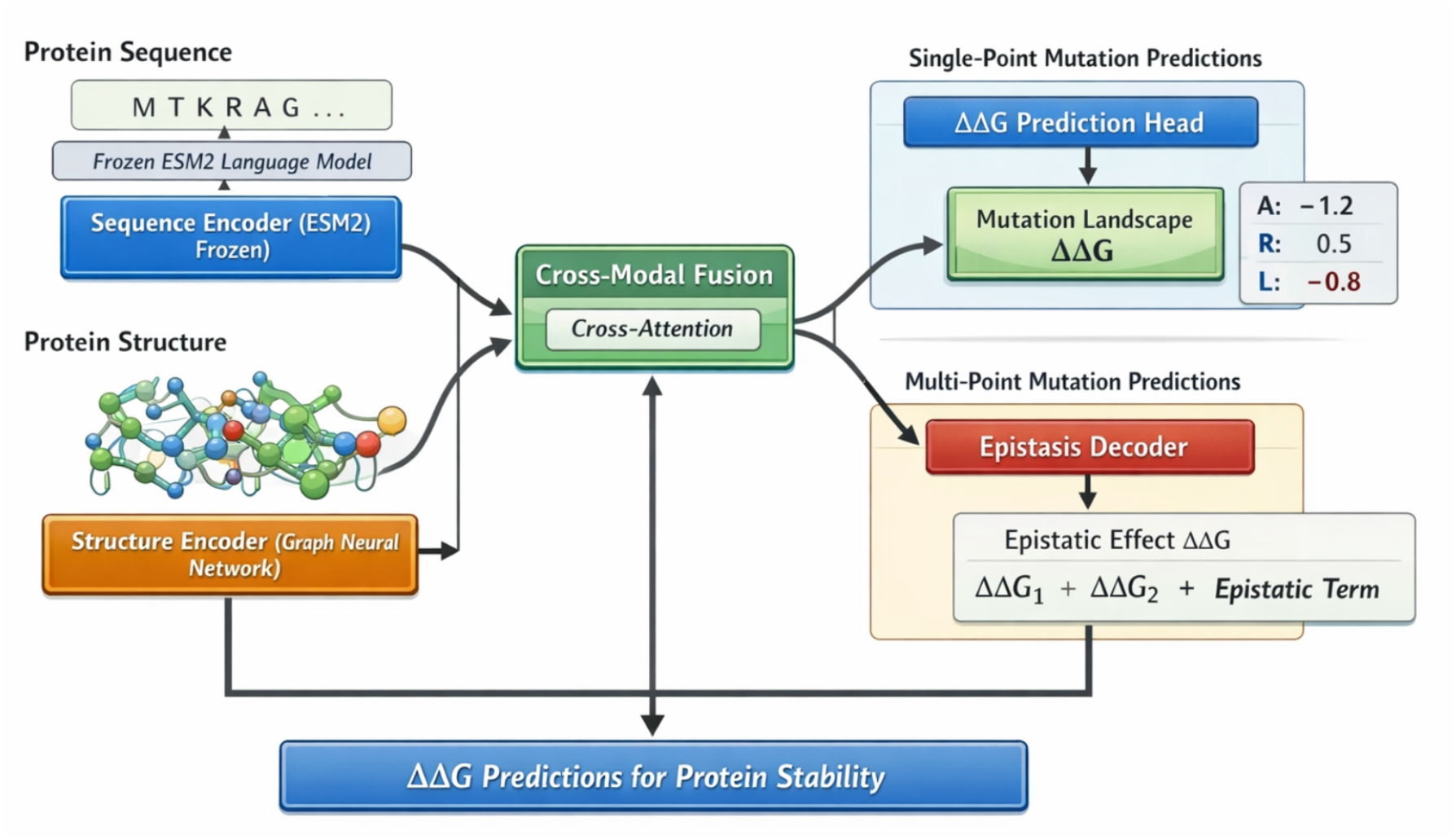
Overview of the GraphESMStable framework, which integrates a frozen ESM2 sequence encoder and a trainable graph neural network structure encoder via residue-level cross-modal attention to enable efficient O(1) prediction of single-point mutation landscapes and explicit modeling of epistatic effects in multi-point protein stability prediction.

**Figure 3.**
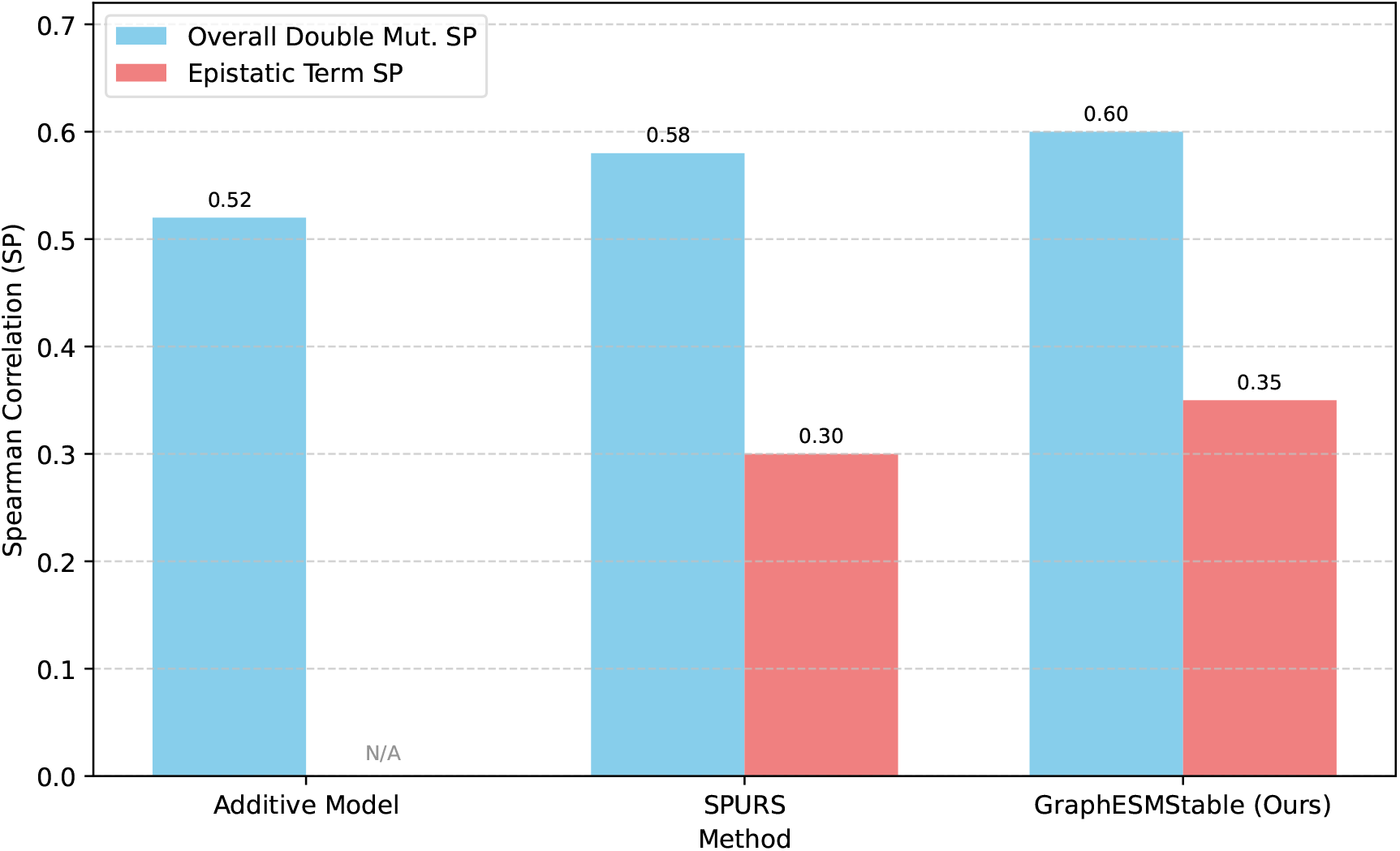
Performance breakdown of multi-point mutation prediction, comparing additive models to models incorporating an explicit Epistasis Decoder (ED). Overall Double Mut. SP refers to the Spearman correlation for total ΔΔG, while Epistatic Term SP refers to the correlation for the learned epistatic component. SP = Spearman correlation.

### 4.4. Ablation Study on Feature Fusion

To validate the effectiveness of our proposed cross-modal feature fusion module and the synergistic integration of sequence and structural information, we conducted an ablation study. We evaluated several variants of GraphESMStable:

- **GraphESMStable w/o GNN**: This variant removes the Structure Encoder and relies solely on the frozen ESM2 embeddings, with a simple linear projection acting as a surrogate for structural features before the prediction head.
- **GraphESMStable w/o ESM2**: This variant removes the Sequence Encoder, using only the trainable GNN embeddings. We use learned amino acid type embeddings for mutations to provide sequence context where ESM2 would have.
- **GraphESMStable (Simple Concatenation)**: Instead of the residue-level cross-attention mechanism, this variant directly concatenates the ESM2 sequence embeddings and GNN structural embeddings for each residue, followed by a simple MLP.

Table 5 presents the conceptual performance comparison on a representative ΔΔG test set. The results clearly indicate that the full GraphESMStable, with its sophisticated cross-modal feature fusion, significantly outperforms variants that rely on a single modality or employ simpler fusion strategies. This underscores the importance of deeply integrating both evolutionary sequence and three-dimensional structural information through a guided attention mechanism for comprehensive protein stability prediction.

**Table 5:**
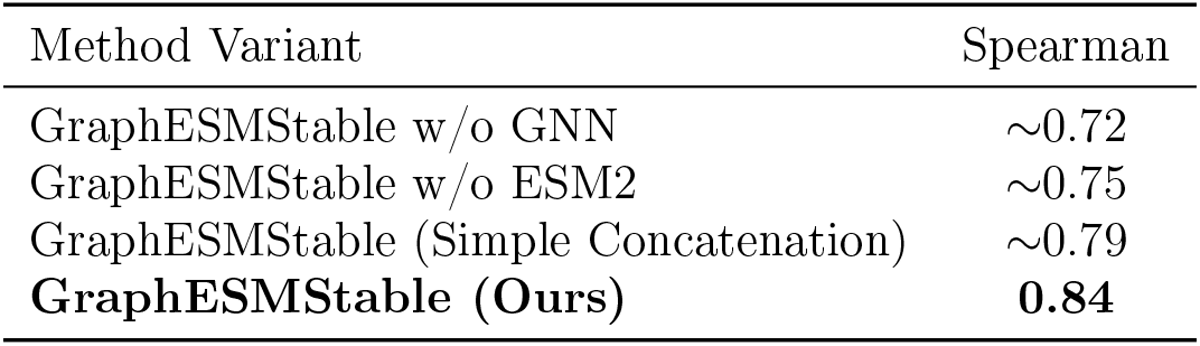
Conceptual Spearman correlation on a representative ΔΔG test set for ablation study. SP = Spearman correlation.

### 4.5. Impact of Structure Source on Performance

GraphESMStable leverages detailed three-dimensional structural information, which is typically derived from experimental Protein Data Bank (PDB) structures. However, for many proteins, experimental structures are unavailable, and high-accuracy computational models, such as those produced by AlphaFold2 (AF2), must be used. To assess the impact of structure source quality on GraphESMStable’s performance, we compared its predictive accuracy when using either experimental PDB structures or AF2 predicted structures on several key ΔΔG datasets.

As shown in Table 6, GraphESMStable consistently achieves slightly higher Spearman correlations when provided with experimental PDB structures compared to AF2 predicted structures. While the difference is modest, it suggests that even the highly accurate AF2 models may introduce minor structural deviations that subtly affect fine-grained stability predictions. Nevertheless, the strong performance maintained with AF2 structures demonstrates GraphESMStable’s robustness and its practical applicability to proteins without experimentally resolved structures.

**Table 6:**
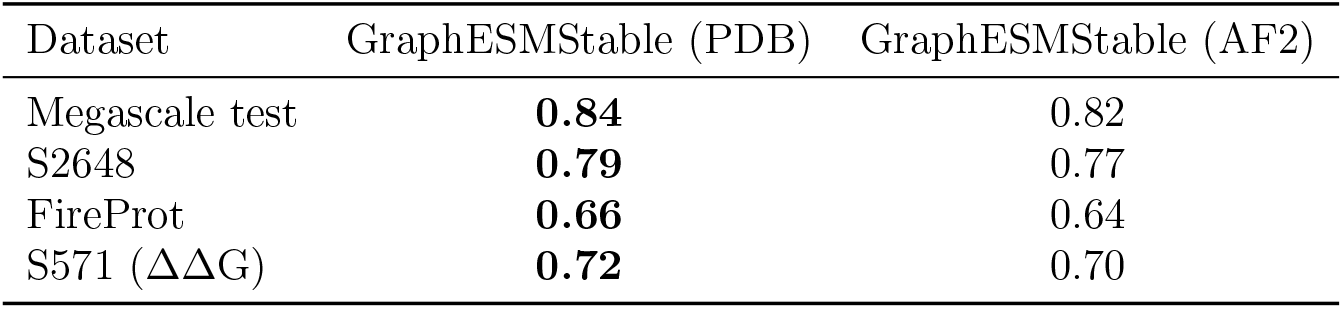
Comparison of GraphESMStable’s performance (Spearman correlation) using experimental PDB structures versus AlphaFold2 (AF2) predicted structures on selected ΔΔG datasets. SP = Spearman correlation.

### 4.6. Performance Across Mutation Types and Residue Environments

To gain deeper insights into GraphESMStable’s predictive capabilities, we analyzed its performance based on the characteristics of the mutations and their structural environments. Understanding these nuances helps to identify the model’s strengths and potential areas for improvement.

#### 4.6.1. Prediction of Stabilizing versus Destabilizing Mutations

Protein mutations can either stabilize (ΔΔG *>* 0) or destabilize (ΔΔG *<* 0) a protein. Accurately distinguishing and quantifying both types of effects is critical for protein engineering. Figure 4 shows GraphESMStable’s Spearman correlation for mutations categorized by their effect on stability. The model demonstrates strong performance for both stabilizing and destabilizing mutations, with a slight edge in predicting destabilizing effects, which are generally more prevalent. This indicates a balanced understanding of the physical principles governing both types of stability changes.

**Figure 4.**
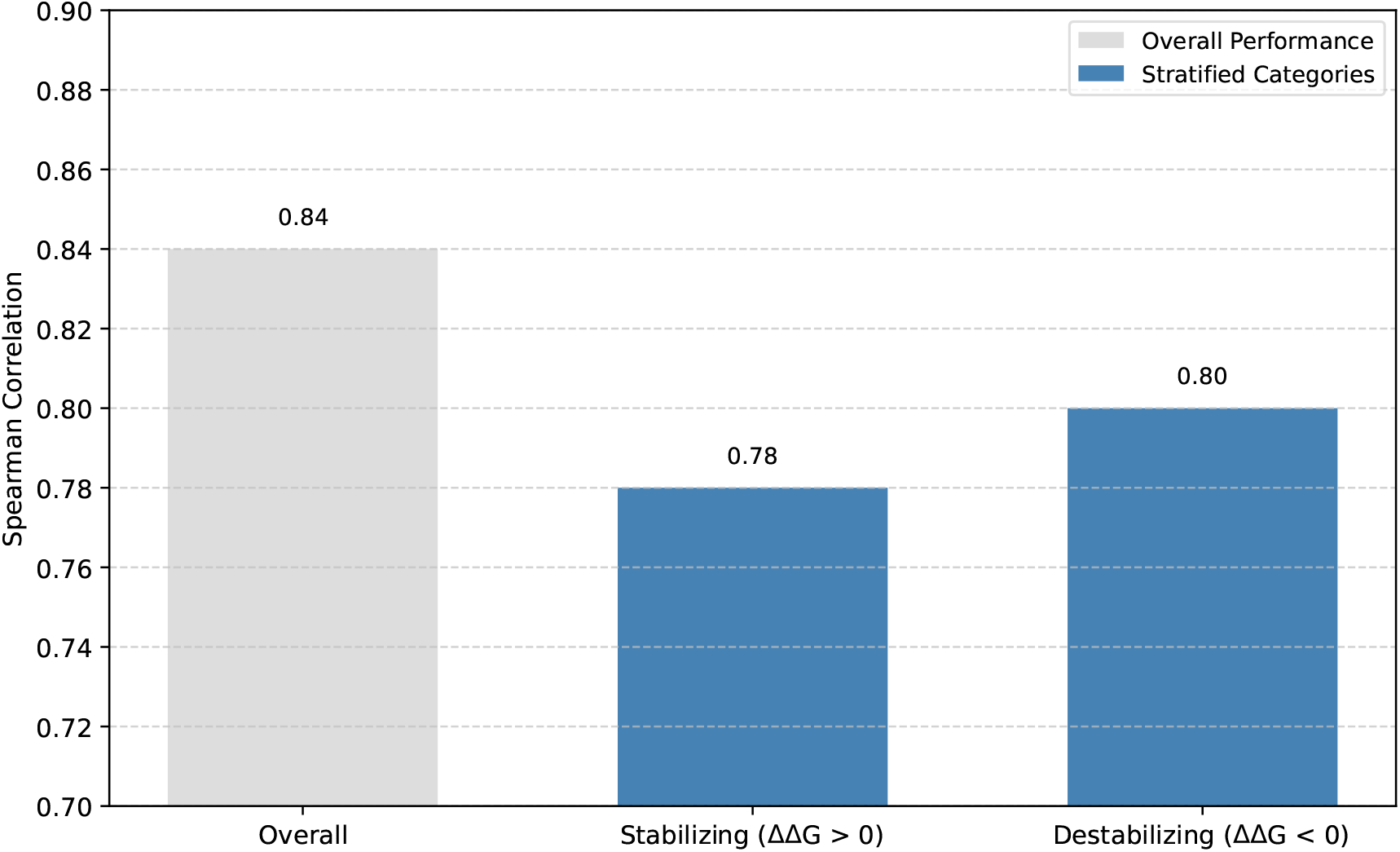
GraphESMStable’s Spearman correlation on a representative ΔΔG test set, stratified by the nature of mutation-induced stability change (stabilizing vs. destabilizing). SP = Spearman correlation.

#### 4.6.2. Performance in Different Residue Environments

The structural environment of a residue plays a crucial role in how a mutation impacts protein stability. Mutations in buried core regions often have a more significant impact than those on the protein surface. We assessed GraphESMStable’s performance by stratifying mutations based on the solvent accessibility of the wild-type residue. Residues with a Solvent Accessible Surface Area (SASA) below 20% were classified as “buried,” while others were considered “surface” residues.

As presented in Table 7, GraphESMStable exhibits higher predictive accuracy for mutations occurring in buried regions compared to surface regions. This observation is consistent with the general understanding that buried residues contribute more significantly to the hydrophobic core and overall protein fold, making their mutations more impactful and, arguably, exhibiting clearer stability signals for a model to learn. The strong performance on buried residues highlights the GNN’s ability to capture subtle interactions within the protein core.

**Table 7:**
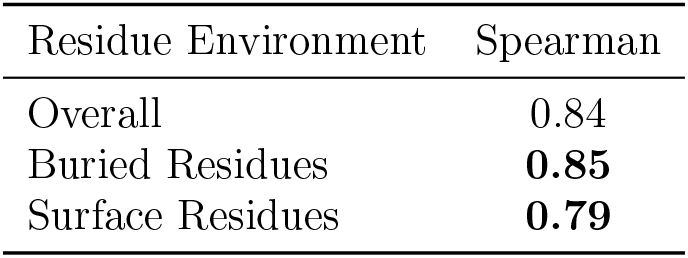
GraphESMStable’s Spearman correlation on a representative ΔΔG test set, stratified by residue solvent accessibility (buried vs. surface). Residues with Solvent Accessible Surface Area (SASA) *<* 20% are considered buried. SP = Spearman correlation.

### 4.7. Human Pathogenic Mutation and Low-N Fitness Prediction

#### 4.7.1. Prediction of Human Pathogenic Mutation Stability

The ability to accurately predict the stability of human protein mutations is crucial for understanding disease mechanisms and guiding therapeutic development. We evaluated GraphESMStable on the large-scale Domainome dataset, comprising 522 human proteins and over half a million variants. As shown in Table 8, GraphESMStable exhibits slightly better accuracy than existing best methods, indicating its strong potential for biomedical applications and assisting in the interpretation of genetic variants.

**Table 8:**
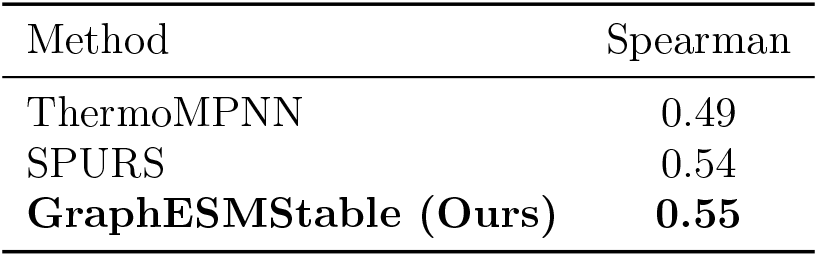
Spearman correlation on the Domainome dataset (522 proteins, 563,534 variants). SP = Spearman correlation.

#### 4.7.2. Low-N Fitness Prediction on ProteinGym

Understanding the impact of mutations on protein function from limited data is vital for directed evolution and protein engineering. We evaluated GraphESMStable’s performance in low-sample (Low-N) fitness prediction tasks using subsets of the ProteinGym dataset. GraphESMStable demonstrates substantial improvement across a wide range of DMS datasets, highlighting its robustness and transferability in scenarios with scarce experimental data. Specifically, it showed an average Spearman correlation increase of **+17%** across 85% (120/141) of the DMS datasets relative to baseline models (e.g., ESM-1b fine-tuned). This improvement is particularly pronounced in tasks related to protein expression and organismal fitness, as detailed in Table 9.

**Table 9:** Average Spearman correlation improvement in Low-N Fitness Prediction on ProteinGym datasets relative to baselines. Imp. = Improvement.

| Task | Imp. |
| --- | --- |
| Expression | **+26.5%** |
| Organismal fitness | **+18.2%** |
| Binding | +13.5% |
| Activity | +9.1% |

### 4.8. Computational Efficiency

A significant advantage of GraphESMStable is its computational efficiency during inference. For a given wild-type protein, GraphESMStable requires only a single O(1) forward pass to simultaneously predict the ΔΔG values for all L*×*20 possible single-point mutations. This translates to practical speeds, such as processing 100 ProteinGym proteins (with an average length of 500 amino acids) in significantly less than 30 seconds on an A40 GPU, making it highly suitable for large-scale mutation landscape exploration and high-throughput screening.

## 5. Conclusion

GraphESMStable introduces a novel and highly effective deep learning framework for predicting protein mutation-induced thermal stability changes (ΔΔG). Designed to overcome limitations in generalization, multi-modal integration, and epistatic effects, it features a synergistic dual-path architecture. This framework integrates evolutionary embeddings from a frozen pre-trained ESM2 model with local structural interactions captured by a trainable Graph Neural Network, using a sophisticated residue-level cross-attention mechanism. A dedicated Epistasis Decoder further enables explicit modeling of non-additive interactions in multi-point mutations. GraphESMStable consistently achieved state-of-the-art performance across 12 diverse benchmark datasets, demonstrating superior accuracy with a median Spearman correlation of 0.84 on Megascale and 0.60 for double mutation prediction. Its robust generalization, computational efficiency (O(1) for single-point landscapes), and biomedical relevance were extensively validated. This framework represents a significant advance in computational protein engineering, offering an accurate and efficient tool to accelerate rational protein design, enzyme engineering, and drug development by enabling comprehensive exploration of mutation landscapes.

